# Brief Apnea and Hypoventilation Reduces Seizure Duration and Shifts Seizure Location for Several Hours in a Model of Severe Traumatic Brain Injury

**DOI:** 10.1101/2023.09.21.558686

**Authors:** Frances Rodriguez Lara, Praneel Sunkavalli, Michael Mikaelian, Bryan Golemb, Kevin Staley, Beth Costine-Bartell

**Affiliations:** Department of Neurosurgery, Massachusetts General Hospital, Boston, MA; Department of Neurosurgery, Harvard Medical School, Boston, MA; Department of Neurology, Massachusetts General Hospital, Boston, MA; Department of Neurology, Harvard Medical School, Boston, MA

**Keywords:** traumatic seizure, status epilepticus, hemorrhage, hypoxic-ischemic injury

## Abstract

**Objective:** Seizures are difficult to control in infants and toddlers. Seizures with periods of apnea and hypoventilation are common following severe traumatic brain injury (TBI). In our multifactorial, severe TBI model (cortical impact, mass effect, subdural hematoma, subarachnoid hemorrhage, seizures induced with kainic acid, and brief apnea and hypoventilation), we observed that brief apnea with hypoventilation (A&H) after induced seizure acutely interrupted seizures, leading us to hypothesize that brief A&H might reduce seizure duration beyond the brief hypoxia and hypercapnia for several hours thereafter. The effects of the timing of A&H on seizure duration and location might inform the pathophysiology of this hypoxic-ischemic injury as well as potential treatments.

**Methods:** Piglets (1 week or 1 month old) received multi-factorial injuries. Apnea and hypoventilation (1 min apnea, 10 min hypoventilation; A&H) was induced either before or after seizure induction, or as a control piglets received subdural/subarachnoid hematoma and seizure without A&H. In an intensive care unit, piglets were sedated, intubated, mechanically ventilated, and epidural EEG was recorded for an average of 18 hours after seizure induction.

**Results:** In our severe TBI model, A&H after seizure reduced ipsilateral seizure burden by 80% compared to the same injuries without A&H. In the A&H before seizure induction group, more piglets had exclusively contralateral seizures though most piglets in all groups had seizures that shifted location throughout the several hours of seizure. After 8-10 hours, seizures transitioned to interictal epileptiform discharges regardless of timing of A&H.

**Significance:** Even brief A&H may alter traumatic seizures We will address the possibility of induced spreading depolarization prior to preclinical investigations of hypercapnia with normoxia, with controlled intracranial pressure, as a therapeutic option for children with status epilepticus after hemorrhagic TBI.

## INTRODUCTION

An escalating anti-epileptic drug protocol is the standard of care in the PICU for the treatment of traumatic seizures. While it is known that worse brain injuries result in more seizures, it is not known if seizures make the tissue damage worse directly driving pathophysiology. Traumatic brain injury (TBI) in infants presenting with an acute subdural hematoma (SDH) is often characterized by seizures with brief periods of apnea and hypoventilation (A&H).^1, 2^ Brief A&H lead to a myriad of effects, one of which is an increase in the concentration of CO_2_ in the blood and tissues. The brain is highly sensitive to small changes in CO_2_ via proton sensitive sodium receptors-acid-sensing ion channel 1a.^3^ Increased CO_2_ during normoxia has been long-characterized to be anti-convulsant in both humans and in animal models.^4-8^ Inhaled medical carbogen (5% CO_2_ and 95% O_2_) rapidly stopped seizures in children with drug-resistant epilepsy.^6, 9, 10^ Medical carbogen has been hypothesized to only have a transient anti-convulsant effect as pre-treatment with carbogen did not prevent the pro-convulsant effect of hyperventilation on children with absence seizures.^6^

Though mechanisms of CO_2_’s anti-convulsant effects are well-established, the effect of timing of A&H on seizure localization and duration in severe TBI has not been studied. In severe TBI, where there is often hemorrhage, seizures, and brief apnea and / or hypoventilation, the apnea and hypoventilation might not be of sufficiently long duration to cause an asphyxial pattern of injury. However, the apnea and hypoventilation might act synergistically with the hemorrhage and/or seizure to trigger metabolic mismatch within the brain tissue.^11^ In young children with subdural hematoma after TBI, if the hypodensity as observed via computed tomography, expands to the majority of the hemisphere (termed hemispheric hypodensity; HH) the rate of morbidity and mortality increases.^12-15^ During the development of our model, we titrated the length of apnea such that apnea did not induce cardiac arrest nor an asphyxiation damage pattern in the brain (bilateral, widespread with particularly susceptibility of damage at the sulci/ capillary watershed areas)^16^ with the goal that A&H would act synergistically with other injuries and insults such that no one injury or insult causes pervasive damage but only in concert together.^16,17^

While inducing the HH model injuries, we observed that 1 minute of apnea paused induced seizures. The EEG was suppressed and the seizure would pause or even stop. Sometimes, an additional dose of kainic acid was required to re-initiate the seizure. Here, we test the hypothesis that brief A&H after seizure induction may alter the course of induced seizure for several hours during normoxia and normocapnia as detected via EEG in our intensive care unit where piglets are sedated, intubated, and mechanically ventilated. Understanding the role of brief A&H on seizures might aid our understanding of the pathophysiology of this condition and may aid in the development of therapeutics to stop the progression of pathophysiology before it evolves into HH.

## METHODS

### Surgery and EEG Recordings

Male and female Yorkshire piglets and were housed and fed as previously described.^16, 18^ The protocols and procedures described were in accordance with the guidelines of the American Veterinary Association and the National Institutes of Health and were approved by the Institutional Animal Care and Use Committee at Massachusetts General Hospital in accordance with US Federal laws. The principles of replacement, reduction, and refinement were considered when assigning animals to this study.

In our model of HH where the multi-factorial injuries and insults that are observed in human cases of HH are induced to one-week (developmentally similar to human infants; “infants”) and one-month (developmentally similar to human toddlers; “toddlers”) piglets and results in widespread hypoxic-ischemic tissue damage beyond the site of focal injuries where the pattern is age-dependent when injuries are scaled to estimated brain volume.^17^

During the period of the study, piglets were assigned to this study in addition to a study currently under review where piglets were randomly assigned to treatment groups; piglets were used from groups that received the full HH model injuries where A&H was *after* seizure, had seizure *without* A&H (with or without subdural/subarachnoid hemorrhage, which served as a benchmark controls (**Table 1**). Additionally, pigs were serially assigned to receive the full HH model injuries with A&H *before* seizure (**Table 1**). Eight piglets were excluded due to animal or equipment problems that prevented the execution of the experiment and/or resulted in less than 7 hours of EEG that could be analyzed from both hemispheres (**Table 1**). A total of 16 piglets were included in the study (**Table 1**). No individual data points were excluded. Those analyzing the EEG were blinded to piglet age and treatment and 60% of the EEG analyses was audited by a second or third reviewer for accuracy.

**Table 1.** Piglets receiving HH model (A&H before or after seizure induced) or seizure without A&H.

| **Total Piglets During the Period (N = 24)** |  |
| --- | --- |
| The overall total during the 3.75-year period that were in the appropriate treatment groups. This period was when intracranial, epidural EEG was typically recorded for several hours. |  |
| **Exclusions (N = 8)** |  |
| <u>EEG recording was less than 7 hours:</u><br>Piglet became extubated during a convulsion and was unable to be resuscitated. (411, A&H after seizure)<br>Piglet was recovered from anesthesia/ the EEG electrodes were pulled. (409, A&H after seizure)<br>Piglet had pre-existing, extensive pneumonia that led to persistent hypercapnia and was euthanized early (449, A&H before seizure). | infants: n = 1 (499)<br>toddlers: n = 2 (409, 411) |
| Isoflurane was administered for several hours to reduce the seizure to treat hypotension so the subject could still survive long enough for the histology study (414, A&H after seizure) | infants: n = 0<br>toddlers: n = 1 |
| The arterial blood gases at the pre-injury timepoint indicated respiratory acidosis likely from a difficult intubation and inadvertent apnea or hypoventilation before injuries. (415; A&H after seizure) | Infants: n = 1<br>toddlers: n = 0 |
| <u>Equipment problems:</u><br>Poor EEG signal on the ipsilateral hemisphere that precluded EEG analysis (447, seizure without apnea).<br>New gas analyzer for exhaled isoflurane was of different sensitivity than previous gas analyzer and the wean off of isoflurane was not successful (466, A&H after seizure). | infants: n = 1 (447)<br>toddlers: n = 2 (410, 466) |
| **Total Included in the Study (N = 16)** |  |
| Full HH model, A&H **after** seizure (n = 8)<br>Randomly assigned | infants: 3<br>toddlers: 5 |
| Full HH model, A&H **before** seizure (n = 5)<br>Serially assigned during the same period | infants: 2<br>toddlers: 3 |
| Benchmark Control:<br>Seizure or Seizure + hemorrhage with no A&H (n = 3)<br>Randomly assigned | infants: 1<br>toddlers: 2 |

The HH injuries were induced under anesthesia and scaled to brain volume as previously described.^16, 17^ Briefly, anesthesia was induced with isoflurane and piglets were intubated and mechanically ventilated so that end tidal CO_2_ was maintained between around 40 mmHg. A craniectomy was performed over the coronal suture and a spring-loaded device was deployed to the rostral gyrus.^19^ Pigs were switched from 100% medical oxygen to compressed room air for cortical impact and thereafter. After cortical impact, piglets were transitioned to a non-isoflurane anesthetic to allow for generation of seizures: dexmedetomidine (5–20 μg/kg/h vs. 10-20 μg/kg/h), morphine (1.75 mg/kg/h “infants” vs. 3.5 mg/kg/h “toddlers”), and boli of rocuronium for muscle relaxation (0.30 -

0.60 mg/kg). While pigs were weaned from isoflurane, a midline shift was induced with a Foley catheter balloon displacing 1% of brain volume for 30 minutes.

After end-tidal isoflurane was 0 for 30 minutes, seizures were induced was with kainic acid (42 - 63 μg/kg) mixed into the blood that was placed into the dura (4.5 mL in “infants”, 5.4 mL in “toddlers”) to create a subdural hemorrhage, which often also results in a subarachnoid hemorrhage. An additional dose of kainic acid was administered if needed to induce seizures.

Apnea and hypoventilation were induced prior to induced seizures, after induced seizures or not at all (**Figure 1**). Piglets, who were already sedated, were paralyzed (0.3 – 0.6 mg/kg rocuronium), and mechanically ventilated and brief apnea (1 minute) was induced by turning off the ventilator and clamping the endotracheal tube. After the end of apnea, respiration was resumed for 1 minute on room air and then hypoventilation/hypercapnia was induced by continuing paralysis, administering 2-6 breaths per minute with room air for 10 minutes to achieve a peak end tidal CO_2_ 60-75 mmHg. Oxygen was mixed with room air to keep peripheral tissue oxygenation above 85% during hypoventilation / hypercapnia.

**Figure 1.**
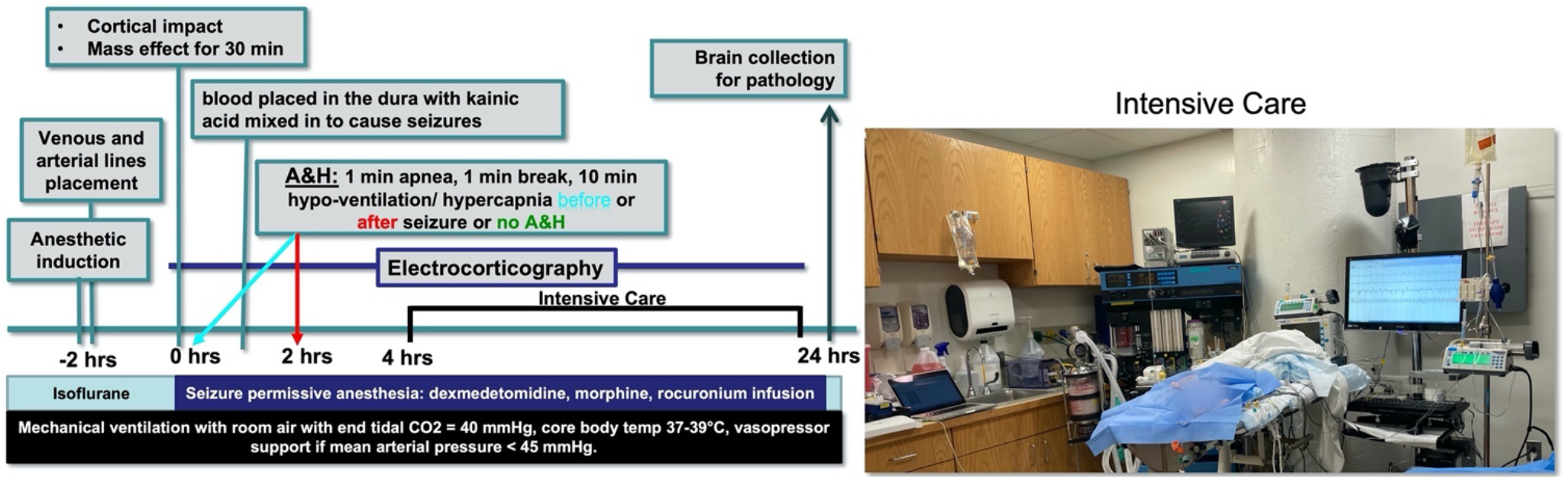
The experimental timeline. Experimental schematic displaying the timing of injuries and insults. A&H (1 minute of apnea, 1 minute break, 10 minutes of hypoventilation/hypercapnia) occurred either prior to kainic acid administration (mixed in with blood injected into the dura), after kainic acid administration or did not occur. Intensive care with ventilator, video EEG monitoring, IV pumps administering seizure permissive anesthesia, and monitoring demonstrated.

Chlorpromazine was administered if needed for additional sedation (0.5–1 mg/kg bolus “infants”; n = 1, 1 - 2 mg/kg, “toddlers”; n = 3). Nitrous oxide (30-50% mixed with 100% oxygen) was administered as needed to achieve comfortable sedation and muscle relaxation in the piglets (n = 4 “toddlers; n = 0 “infants”). If the pulse oximeter was not functioning due to poor peripheral circulation (from poor cardiac function), then the room air was mixed with 100% oxygen. Additionally, piglets were transported on 100% isoflurane to perfusion in those where the brain was collected for histological analysis.

EEG was recorded as previously described using electrode strips containing 4 electrodes on each hemisphere epidurally (AdTech) creating a 4-channel bipolar montage resulting in 2 channels per hemisphere (**Figure 2A**).^20^ A ground electrode was placed on the piglet’s leg and a reference electrode under the scalp. Video EEG was recorded on an Natus XL-Tech EEG unit (amplifier: ref 10388; amplifier break out box: ref 012378) with a 60 Hz Notch filter recorded in NeuroWorks (Natus Neurology) prior to application of injuries and recording continuously (except for 15 - 30 minutes during transport from the operating room to the intensive care unit) for up to 25 hrs after injury while animals were sedated, intubated, and mechanically ventilated in an intensive care unit.

**Figure 2.**
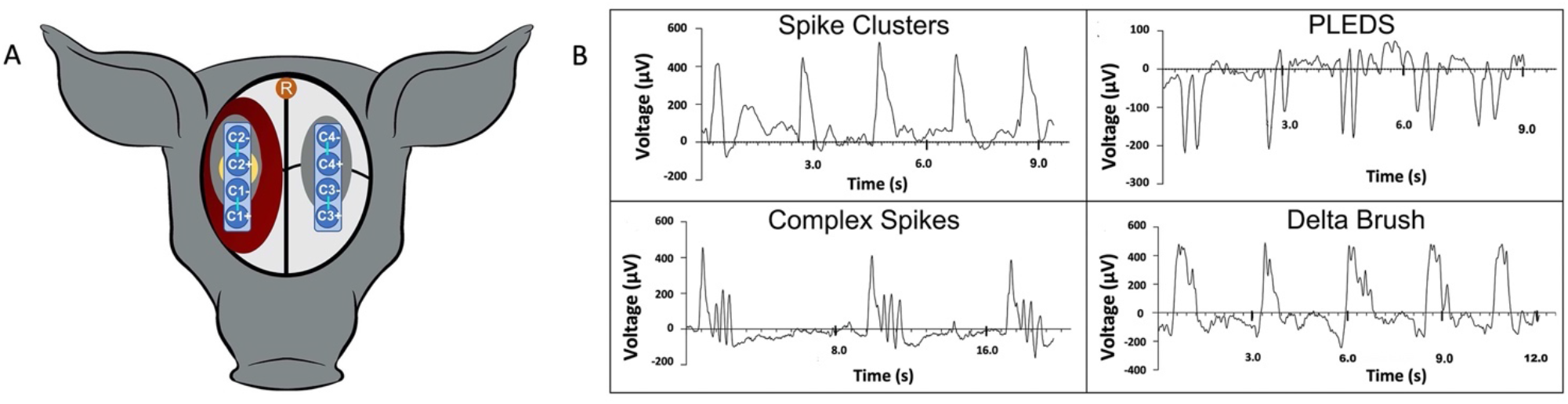
EEG montage and example of interictal epileptiform discharges. **A**. 4-channel bipolar montage with electrode strips placed epidurally centered over the coronal suture with 2 channels per hemisphere (Channel 1 = C1, Channel 2 = C2 etc.; gray: burr hole, yellow: site of cortical impact, red: subdural blood placement, electrode per channel connected via turquoise line). A reference electrode (orange, R) was placed under the scalp. A ground electrode was placed on the piglet’s leg. **B**. Example of spike clusters, periodic lateralized epileptiform discharges (PLEDS), complex spikes, and delta brushes. PLEDS were defined as spikes at a lower frequency than seizures (20 - 100 ms in duration, 1.0 - 1.9 Hz). Spike clusters were low frequency clusters of spikes (20 - 100 ms in duration, 0.15 - 0.6 Hz)^34^. Slow waves with multiple spike inflections on the top of the wave were categorized as delta brushes (200-300 ms in duration, 1.0 - 1.7 Hz)^35^. Complex spikes were defined as paroxysmal discharges distinct from background activity (> 75 μV amplitude off baseline) with variable spike amplitudes of variable duration and variable frequency (0.15 Hz - 48 Hz).

The EEG was analyzed for IED’s, seizures, and suppression during apnea (**Figure 2, 3**). EDF files were imported into NeuroScore (Data Science International) and a Finite Impulse Response filter with 0.5 - 55 Hz frequency was used to reduce noise. A spike was defined as a transient discharge with a duration of 20-100 ms with an amplitude deflecting ≥ approximately 100 μV off baseline. Baseline was estimated by finding the average amplitude of waves in the recording prior to any injuries or insults. Seizures were defined as spikes with frequency of 2 Hz for at least 10 seconds. Seizures were categorized into ipsilateral seizures (ipsilateral to focal injuries), contralateral seizures (hemisphere opposite to focal injuries), or generalized seizures (at least one channel in both hemispheres). Specific types of IED’s that were quantified included periodic lateral discharges (PLEDS), spike clusters, delta brush, and complex spikes (**Figure 2B**). IED’s or seizures were marked if present on one or more channels. If a seizure was present on one or more channels along with IED’s on other channels, IED’s at this time were not quantified; this was a rare event. Physiologic waves such k-complexes were observed but not quantified as an IED. Artifact was considered any large amplitude discharges if the event has a constant amplitude throughout the discharge (complex spike but constant amplitude) and any waveforms over 48 Hz (muscle artifact). Here, seizure duration was not different between ages (P = 0.73), and therefore, ages were combined for analysis. EEG suppression during apnea was quantified when the range of EEG was an amplitude within 50 mV and -50 mV in amplitude during induced apnea until resolved.

**Figure 3.**
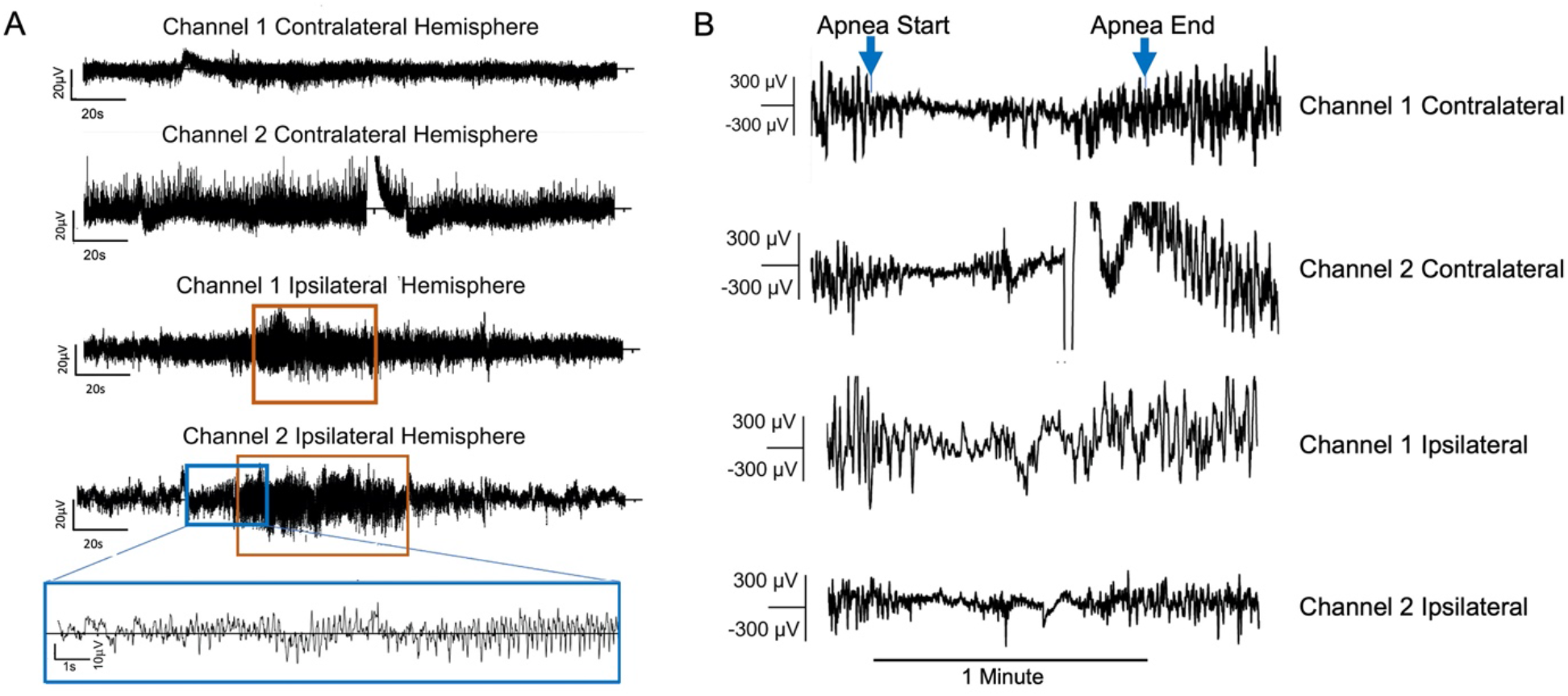
Examples of a seizure and EEG suppression during apnea. **A**. The 4-channel bipolar montage with the top two channels of the contralateral hemisphere displaying baseline activity and a seizure (orange box) is detected on both channels of the ipsilateral hemisphere. The blue box displays the expanded timescale of the initiation of the seizure where amplitude and frequency increased from baseline and evolves into a seizure. **B**. One minute of apnea resulted in EEG suppression during apnea.

### 2.2 Physiology and Blood Collection and Statistical Analysis

Arterial blood was collected once prior to the injury and then at intervals of 1-, 3-, 6-, 8-, 12-, 16, and 24-hours post-injury and analyzed with an I-STAT Handheld device (Abbot) to measure PO_2_, PCO_2_, base excess in the extracellular fluid compartment, glucose, pH, and lactate. If glucose was < 70 mg/dL, then 5% dextrose was administered until remediated. At 24 hours post-injury, piglets were deeply anesthetized with 3-5% isoflurane and perfused and exsanguinated or piglets were euthanized with Euthasol (Virbac AH, Inc.) for fresh brain collection for another study (under review).

Data are presented as means ± standard deviation. Statistical analyses were pre-planned. The main effects of treatment (A&H before seizure, A&H after seizure, or no A&H) and time and the interaction on seizure duration, total IED’s, and blood gases and metabolites were tested via a two-way ANOVA followed by Tukey’s post hoc tests. The main effects of treatment and total seizure duration/location and the interaction was tested via a using a two-way ANOVA followed by Tukey’s post hoc tests. Because a striking pattern on age and IED type was observed, the main effect of IED type (delta brush or complex spikes) and age and the interaction were tested via a two-way ANOVA. The effect of treatment on total IED’s or time of last ipsilateral seizure after seizure induction were tested with a one-way ANOVA. The effect of A&H on end tidal CO_2_ and oxygen saturation were tested via a two-way ANOVA followed by Tukey’s post-hoc tests. The dose of kainic acid required to induce seizure between groups (A&H vs. no A&H) and length of EEG suppression during apnea (A&H before seizure vs. A&H after seizure) were tested via an unpaired, two-tailed Student’s T-test. The percentage of piglets with exclusively ipsilateral seizure, exclusively contralateral seizure or alternating among all categories among groups was tested with a Chi Square test on a 3 x 3 contingency table. The number of piglets requiring vasopressors, chest compressions or had an early death due to cardiac issues was compared via Fisher Exact Tests.

## RESULTS

Piglets receiving A&H after seizure exhibited an 80% reduction in the total time spent in seizure than piglets without A&H (9.5 ± 5.7 hrs to 2.5 ± 2.4 hrs, P = 0.002; **Figure 4A, 4B**; main effect of treatment; P = 0.001, a main effect of seizure location, P < 0.000, significant interaction, P = 0.001). Additionally, there was a main effect treatment on time spent in seizure over the ∼20-hour recording period (P = 0.02; repeated measures ANOVA; **Figure 4B**). Piglets were recorded with video EEG for an average of 17.8 ± 4.2 hours. The time of recording did not differ among groups. The length of EEG suppression either A&H before or after the EEG was not different (23.86 ± 22.34 vs. 14.40 ± 12.66 seconds, P = 0.42).

**Figure 4.**
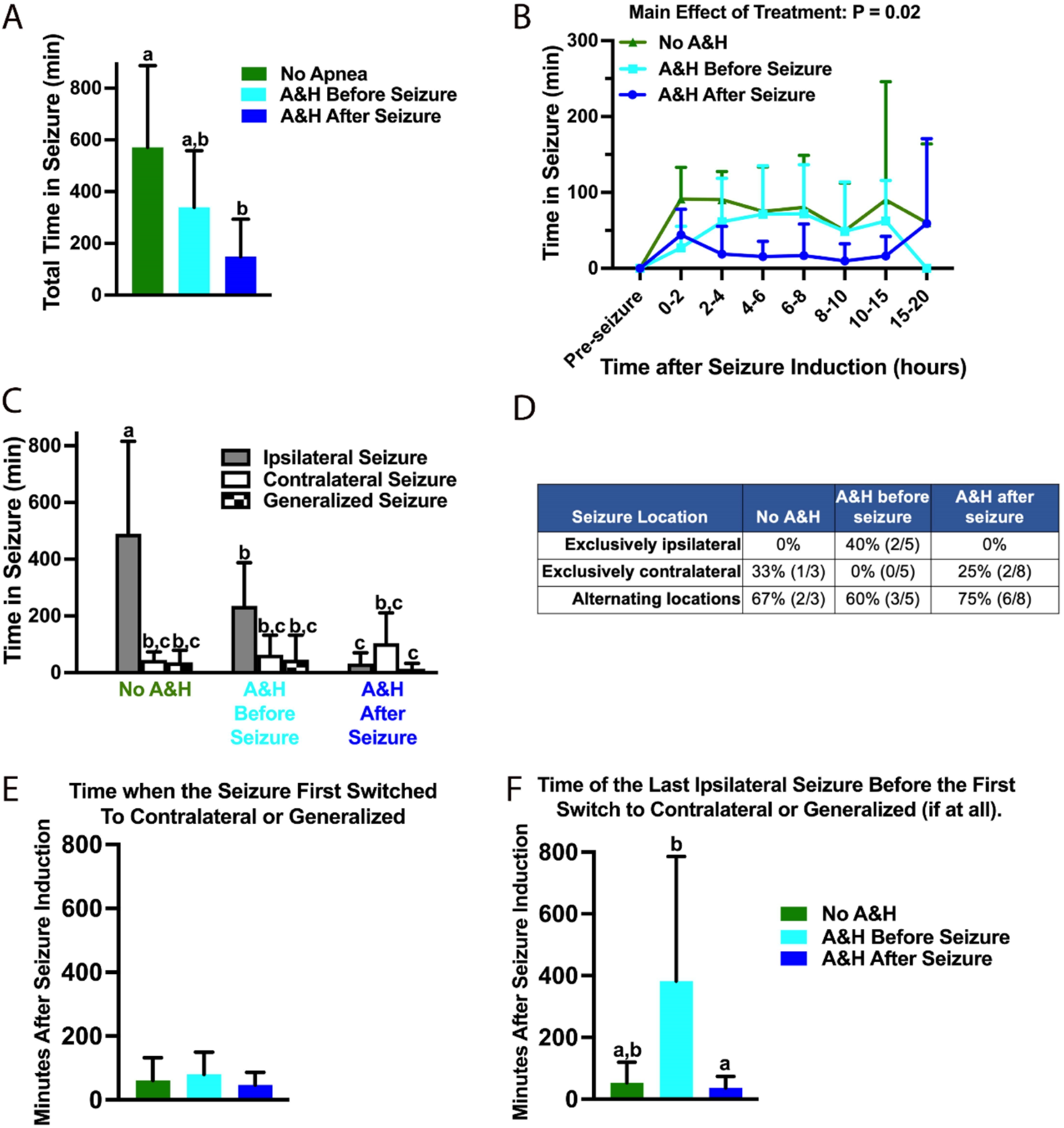
Brief apnea and hypoventilation (A&H) reduced seizure duration and changed the location of the seizure for several hours. **A, B**. The total time spent in seizure was reduced in those receiving A&H after seizure induction (N = 8) vs. those without A&H (N = 3; main effect of treatment, P = 0.001). **C**. The location of the seizure changed with apnea (main effect, P < 0.0001). Without (A&H), seizures were mainly restricted to the hemisphere ipsilateral to where the blood mixed with kainic acid was injected into the subdural space. Apnea before (N = 5) or after seizure (N = 8) resulted in seizures being equivalently distributed between hemisphere or were generalized. ^a,b,c^Means ± SD with different letters differ, P ≤ 0.05. **D**. The percentage of pigs with different locations of seizure: exclusively ipsilateral, exclusively contralateral, or alternating among all categories differed due to A&H (Chi Square, 3 x 3 table, P < 0.0001). To determine if there was a “switch” from ipsilateral to contralateral after A&H, the time after seizure induction of the first contralateral/ generalized (**E**) and the last ipsilateral seizure (**F**) before a switch was noted for each pig. **E**. The time of the first seizure that was contralateral or generalized was not different among treatments. **F**. The time of the last ipsilateral seizure before switch was greater in piglets with A&H before seizure likely due to the piglets that had exclusively ipsilateral seizures. ^a,b^Means ± SD with different letters differ, P < 0.05.

Brief A&H changed the location of the seizure. Seizures were induced with kainic acid mixed with blood placed injected into the dura over the right hemisphere (ipsilateral hemisphere; same location as the cortical impact and induced mass effect). Without A&H, the average time spent in seizure among piglets was greater in the ipsilateral vs. contralateral hemisphere or generalized (**Figure 4C**). Brief A&H before or after seizure induction resulted in the seizure being distributed equally among each hemisphere or were generalized; therefore, the time in seizure in the ipsilateral hemisphere was what was reduced with A&H (**Figure 4C**).

Brief A&H also altered the location of seizure when examining patterns in individual piglets. We observed that among all groups, most piglets had seizures that alternated location during the several hours of seizure (60-75%, **Figure 4D**). However, there was an effect of A&H (P < 0.001, Chi Square) on the percentage of piglets with seizures that were exclusively ipsilateral or contralateral (**Figure 4D**). In piglets exposed to brief A&H before seizure, 40% had seizures exclusive to the ipsilateral hemisphere and none had exclusively contralateral seizures. In contrast piglets with A&H after seizure, none had exclusively ipsilateral seizures and 25% had seizures exclusive to the contralateral hemisphere (**Figure 4D**).

To query if there was a “switch” where seizures were initiated in the ipsilateral hemisphere but then switched the contralateral hemisphere with A&H, we measured the time after seizure induction to the time of the first switch to contralateral hemisphere or to generalized seizure as well as the time of the last ipsilateral seizure. The switch to the first contralateral or generalized seizure was the same among groups and happened within about an hour after induction regardless of exposure or timing of A&H (**Figure 4E**). The timing of the last ipsilateral seizure was only different among groups not due a switch but because a proportion of piglet receiving A&H before had seizure that never left the ipsilateral hemisphere (**Figure 4D**,**F**). Most subjects among all groups had seizures that switched locations regularly over the several hours while A&H before seizure might have a priming effect increasing the rate of exclusive ipsilateral seizure in this group. The difference observed in location in the pigs without A&H and the pigs with A&H after seizure (**Figure 4C**) was the cumulative location for several hours with a reduction in seizure in the ipsilateral hemisphere.

The type of IED differed with age with infant piglets displaying more delta brush IED and toddlers displaying more complex spikes after severe TBI injuries (age x IED interaction; P = 0.05; **Figure 5A**). Treatment did not affect the time spent in IED (**Figure 5B**) but IED’s increased 10 - 15-hour post-seizure induction among all groups (main effect of time, P = 0.01) coincident with the end of seizure activity (**Figure 5C**).

**Figure 5.**
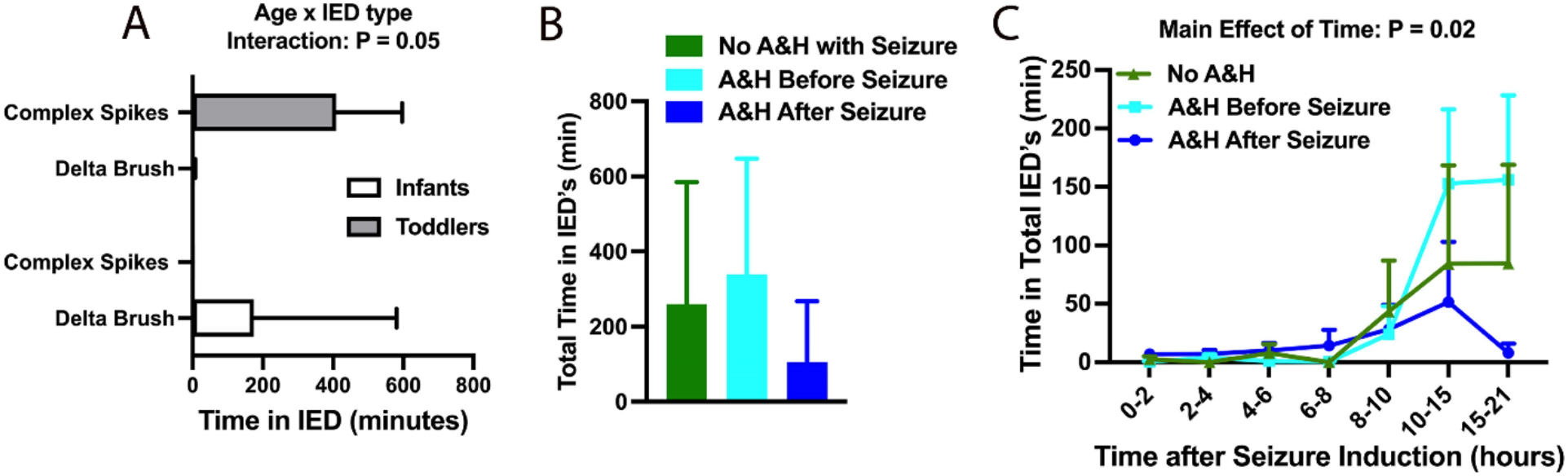
The type of interictal epileptiform discharges (IED’s) was age dependent and increased over time. **A**. There was an interaction with age and IED type with infants having more delta brush and toddlers displaying more complex spikes in this model of severe TBI (two-way ANOVA). There was no effect of brief A&H on the total time spent in IED’s (**B**.), but all groups increased over time (**C**. Main effect of time, P = 0.02, two-way ANOVA).

To examine a protentional indicator of seizure threshold in relation to the timing of A&H, we analyzed the dose of kainic acid that administered required to induce seizure. If seizure was not successful after the initial dose of kainic acid, additional doses were administered into the brain parenchyma to effect. The dose of kainic acid required to initiate seizure was not different among groups.

CO_2_ and oxygen were compared between groups with A&H during the brief A&H via end tidal CO_2_ and oxygen saturation via a pulse oximeter. As expected, hypoventilation caused hypercapnia in both groups (**Figure 6A**). Elevated end tidal CO_2_ resolved within 15 minutes with mechanical ventilation turned on back to the pre-hypoventilation rate. One minute of apnea reduced oxygen saturation and remained lower than pre-apnea in during the hypoventilation in both groups (**Figure 6B**). Oxygen saturation was not different at the 10-minute timepoint during hypoventilation than pre-injury in the A&H before seizure group and took longer to recover in the A&H after seizure group resolving by the next recording (15 minutes after the end of hypoventilation; **Figure 6B**).

**Figure 6.**
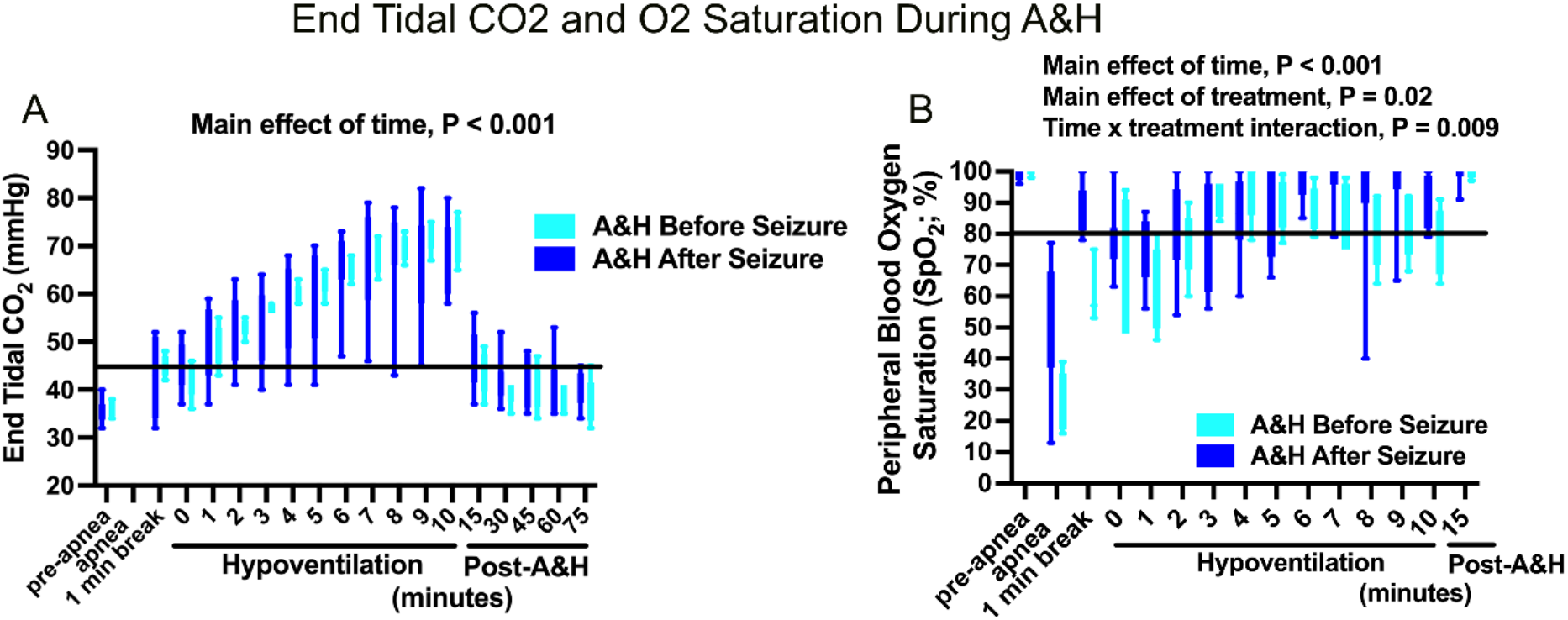
Peripheral oxygen and end tidal CO_2_ during A&H. **A**. As expected, 10 minutes of hypoventilation increased end tidal CO_2_ and was not different between groups with apnea indicating that changes in seizure duration or location for several hours was not due to differences in the induced A&H between groups. **B**. Brief A&H reduced peripheral oxygen saturation both during the 1-minute apnea and in the period of 10 minutes hypoventilation. Despite, mixing 100% oxygen in with room air during hypoventilation, piglets were hypoxemic in at least one of the two groups (below 80% peripheral oxygen) for 8 of the 10 minutes hypoventilation (peripheral oxygen saturation above 80%). Peripheral oxygen saturation was back to pre-A&H concentrations the first time it was measured after A&H. Blood gases were not obtained during brief A&H.

The differences in seizure length cannot be explained by differences in blood gases and metabolites for the duration of the experiment outside of the period of brief A&H. Blood gases were obtained pre-injury and throughout the duration of the experiment. Though there was a main of effect of time on nadir pH, nadir base excess of the extracellular fluid, peak glucose, and peak lactate from pre-injury, 1-2 hrs post-A&H (or similar time), and the peak or nadir, these values did not differ between groups where seizure were induced (**Figure 7)**. Despite the expected changes in metabolites due to the severe TBI injuries and seizures, there was no different in pO_2_ (**Figure 7E**) nor pCO_2_ (**Figure 7D**) during the duration of experiment indicating that the changes in CO_2_ and O_2_ induced here was likely limited during the brief A&H period.

**Figure 7.**
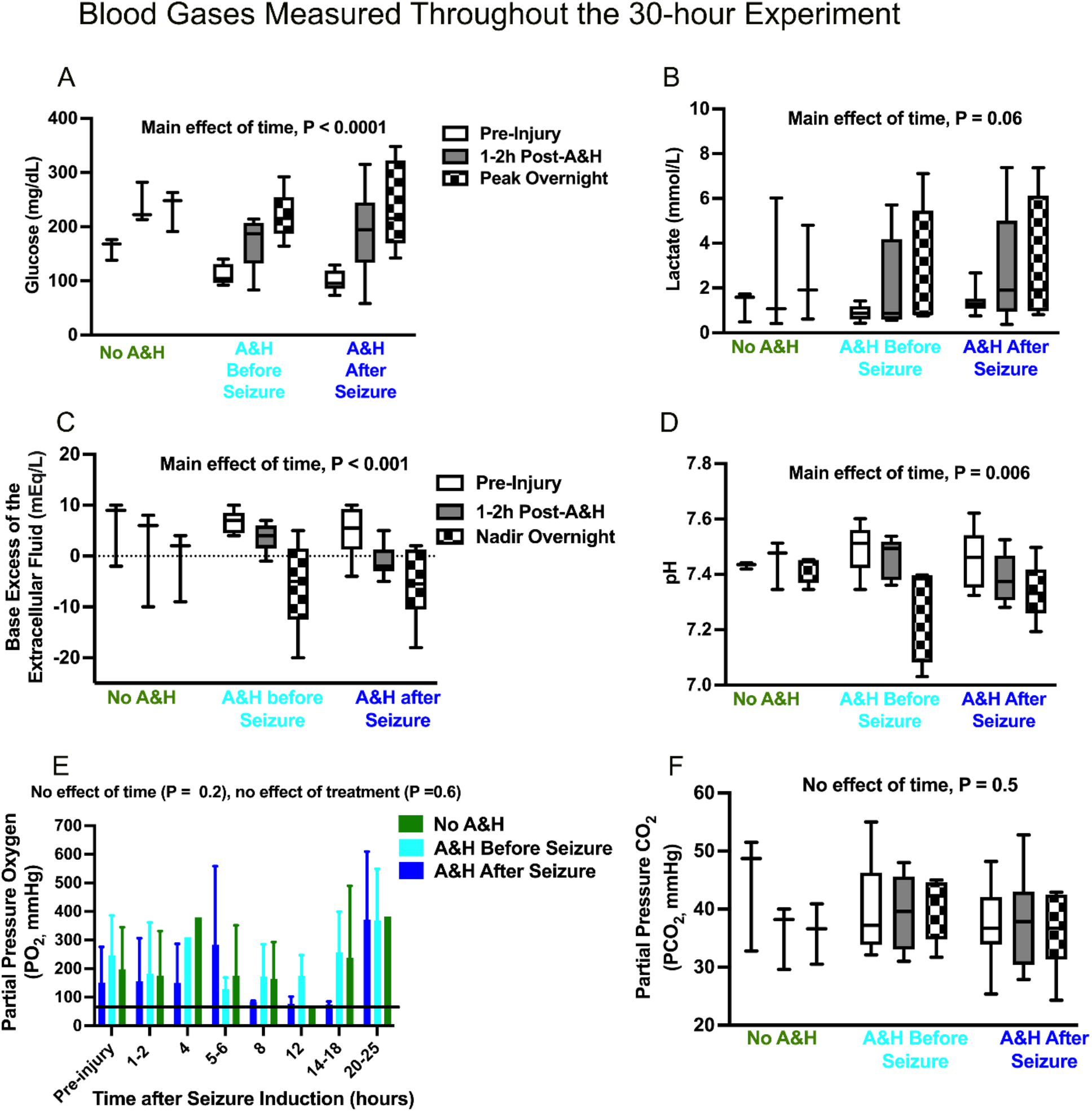
Differences in seizure length were not due to ongoing differences in partial pressure of O_2_ nor CO_2_ after the brief A&H. Severe TBI injuries induced metabolic acidosis while partial pressure oxygen and CO_2_ remain the same over time and among groups. Blood gases and chemistries were measured pre-injury, 1-2 hours post-seizure initiation, and regular intervals for the next 25 hours. Glucose (**A**) and lactate (**B**) increased while base excess of the extracellular fluid (**C**), pH (**D**) all decreased over time (main effect of time) while partial pressure O_2_ (**E**) and CO_2_ (**F**) did not decline over time indicating metabolic acidosis and not respiratory acidosis for the several hours following the brief A&H. High pO_2_ over 100% is due to piglets being ventilated with 100% oxygen at times. Differences in parameters were tested via a two-way ANOVA followed by Tukey’s post-hoc tests. ^a,b^Means ± SD differ, P < 0.05.

If piglets had cardiac complications, it was several hours after the brief A&H and not during A&H. Previous work titrated the duration of apnea so that it did not cause cardiac arrest in these ages of piglets. The number of pigs receiving vasopressors (P = 0.27), requiring chest compressions (P = 0.27) nor dying early due to cardiac dysfunction (P = 0.52) was not different indicating that clinical severity among groups was not different (**Table 2**).

**Table 2.** Rate of administration of vasopressors, chest compressions, and early death due to cardiac complications in piglets assigned to the study (includes excluded pigs with the exception of the piglet with pre-existing pneumonia).

| **Treatment** | **Required Vasopressors** | **Required Chest Compressions** | **Early Death due to primarily due seizures causing cardiac dysfunction** |
| --- | --- | --- | --- |
| No A&H | 25%, 1 (447) of 4 | 0%, 0 of 4 | 0%, 0 of 4 |
| A&H before seizure | 0%, 0 of 5 | 0%, 0 of 5 | 0 %, 0 of 5 |
| A&H after seizure | 30%, 4 (409, 411, 415, 432) of 13 | 30%, 4 (411, 414, 432, 465) of 13 | 23%, 3 (432, 465, 506) of 13 |

## DISCUSSION

Brief A&H shortened seizure duration and dispersed the seizure location in our large animal model of severe TBI. The effect of A&H reducing seizure might be due to the hypercapnia, which decreases pH in the brain parenchyma^21^. Inhaling 10% CO_2_ caused a decline in brain parenchymal pH in seconds^22^. The brain is highly sensitive to changes in pH in order to maintain respiratory homeostasis and pH homeostasis within the brain^23^. Brain pH can be altered by both peripheral pH changes as well as local pH shifts due to neuronal activity and is regulated by an array of mechanisms.^21, 23^ Decreased pH is detected by the acid-sensing ion channel 1a (ASIC1a)^22^. Inhaled 10% CO_2_ terminated pentylenetetrazol-induced seizures in wild-type mice but had no effect in ASIC1a-/-mice.^24^ Additionally, hypercapnia and a fall in pH increases extracellular adenosine concentrations, which inhibits neuronal excitatory activity via adenosine A_1_ receptor^21^. Indeed, the activity of most ion channels is modulated in some way by extra neuronal proton concentration^25, 26^.

Neuronal activity in response to pH extrapolates into a powerful effect on seizures. Hyperventilation (hypocapnia) can stimulate seizures in children with absence epilepsy^6, 27^. Inhaled CO_2_ with normoxia can rapidly terminate seizures and epileptiform activity in patients with drug resistant epilepsy^9^ and can terminate absence seizures initiated by hyperventilation^6^. Hypoventilation in awake patients is highly anxiogenic but exposure to 5% carbogen was well-tolerated for the 20-80 seconds required to stop the seizure^6^.

Although peripheral pH and pCO_2_ during brief A&H were not measured and did not differ for several hours, piglets with longer seizures might have certainly had a lower pH in the brain parenchyma and/or lower intraneuronal pH due to increased activity during seizures, which were not measured here. However, activity-dependent intraneuronal acidification serves as a seizure termination mechanism (via ASIC1a)^22^ though intracellular pH quickly recovers between seizures, an increase in seizure threshold might persist longer^28^.

Alternatively, the hypoxia which was extreme for 1 minute and mild for several additional minutes, might have caused the change in seizure duration without the effect of hypercapnia or in combination with hypercapnia. We are not aware of any literature demonstrating brief A&H, brief apnea alone, brief hypercapnia alone that that suppresses seizures for hours later. However, breathing 5% CO_2_ with normoxia is sufficient for the anti-convulsant effect in epileptics while exposed to 5% CO_2_, but it can be argued that in our model, the persistent effect is due to tissue changes induced by brief hypoxia or hypercapnia or both. Brief A&H superimposed upon seizure induction, cortical impact, midline shift, and subdural hematoma affected the tissue such that seizure generation was promoted for several hours later in the midst of normoxia and normal end tidal and CO_2_. Indeed, although seizure and hemorrhage were induced on the ipsilateral hemisphere and stayed in the ipsilateral hemisphere, in the groups where seizure was induced without A&H or after A&H, when A&H was induced after seizure, the seizure shifted to the contralateral hemisphere. We do not know why this occurred, but brief A&H after the induced seizure might deplete metabolic substrate quickly in the hemisphere where the seizure was induced where A&H is synergistic with the seizure while reserving more substrate in the contralateral hemisphere allowing seizure to continue in that location. Brief A&H during seizure might change the tissue such that tissue threshold for seizure is increased for several hours.

Alternatively, hypoxia in the setting of seizures may induce spreading depression in the ipsilateral hemisphere that had been exposed to kainate^29^. Spreading depression is known to occur after seizures^29^ and after subarachnoid hemorrhage^30^, though the precise mechanisms, induction conditions, and the effect on pathophysiology are still being investigated^31^. Similar to seizure and hypercapnia, spreading depression lowers brain pH and could result in the pattern of reduced ipsilateral seizures. Because spreading depression is usually confined to one hemisphere, it would not have spread to the contralateral hemisphere. This is consistent with our finding that contralateral seizures were not affected by A&H after seizures.

The role of traumatic seizures in the development of hypoxic-ischemic injury is not well understood. While worse brain injuries cause more seizures, it’s not yet understood if seizures play into the pathophysiology of infants and toddlers. Less is understood about if brief hypercapnia (with carbogen) would have a net benefit on tissue damage in the context of subdural/subarachnoid hemorrhage and traumatic seizures in the developing brain. Future directions for this research would be comparing the histopathology of hypoxic-ischemic tissue damage^17, 32^ in the brains of pigs that underwent A&H before or after seizures. A&H might not be necessary for the development of HH as HH has been observed to develop in witnessed accidental cases of TBI where A&H was noted to be absent.^33^ The hypothesis that the reduction of seizures and a shift in location reduces the amount of tissue damage could be tested. Wide band recordings with an extensive EEG montage to test for induction of spreading depression in the large piglet brain will also be very useful. Future work would also aim to test the effect of hypoventilation alone or administration of carbogen with normoxia and compare with A&H to determine if both are needed for the seizure suppressive effect. Inducing brief apnea in infants with a lower oxygen reserve and precarious cardiac condition could do more harm than benefit but administering carbogen has already demonstrated to be safe. Administering carbogen or inducing hypercapnia with 100% oxygen to young children already intubated and sedated would be logistically feasible. Examining the potential anti-seizure effect of hypercapnia for clinical translatability would also require control of intracranial pressure. Seizures are difficult to control in young children where GABA is still depolarizing in brain regions. Mild and brief hypercapnia might be more helpful than GABA agonists in controlling seizures in children, even in the context of severe TBI.

## LIMITATIONS

A limitation of this study is that one treatment group was not randomized. The other treatment groups were randomized for other studies not reported here. However, it was within the same time period as the other, randomized groups as this was our first attempt testing the hypothesis that brief A&H would alter seizure duration and location for several hours. An additional limitation is that our control group. We chose existing groups from other studies not reported here to form our control benchmark group: seizure alone or seizure & subdural hematoma. This group not only did not receive A&H but also did not receive cortical impact and some within the group did not receive subdural hematoma, which might have introduced variation within the group. Blood gases were not measured during the period of brief A&H but only every few hours for the course of the 20-hour experiment. End tidal CO_2_ and pulse oximetry is less reliable than pCO_2_ and pO_2_. Work should continue to test is this phenomenon to determine if it is repeatable in a prospective fully randomized trial with a uniform control group. Pilot data such as these in large animal models is key in order to acquire full funding to test our hypothesis that the reducing seizure and location with brief apnea or brief hypercapnia, while controlling intracranial pressure, will reduce tissue damage in this severe TBI model.

## CONCLUSIONS

Brief apnea (1 minute) followed by hypoventilation (10 minutes) reduced the duration of seizures for several hours if administered before or after seizure induction by reducing the amount of time of seizure in the ipsilateral hemisphere for several hours. Little is known about the interplay of seizures and apnea and hypoventilation severe TBI, which includes subarachnoid and subdural hemorrhage. Future studies will aim to elucidate if a brief period of hypercapnia with controlled intracranial pressure will reduce seizure and the extent of the spread of hypoxic-ischemic tissue damage. Seizures are difficult to control in young children where GABA is still depolarizing in brain regions. Mild and brief hypercapnia with normoxia might be more helpful than GABA agonists in controlling seizures in children, even in the context of severe TBI.

## Abbreviations

A&H: brief apnea and hypoventilation
HH: hemispheric hypodensity
IED: interictal epileptiform discharges
TBI: traumatic brain injury

## AUTHOR DISCLOSURES

None of the authors have conflicts of interest to disclose. The work described here is consistent with the Journal’s guidelines for ethical publication.

## DATA AVAILABILITY STATEMENT

The data that supported the findings of this study are available from the corresponding author upon reasonable request.

## ORICD

*Beth Costine-Bartell* https://orcid.org/0000-0001-7559-9378

## Notes

### Competing Interest Statement

The authors have declared no competing interest.

## REFERENCES

1. Kemp AM, Stoodley N, Cobley C, Coles L, Kemp KW. Apnoea and brain swelling in non-accidental head injury Arch Dis Child. 2003 Jun;88:472–476; discussion 472-476.

2. Ichord RN, Naim M, Pollock AN, Nance ML, Margulies SS, Christian CW. Hypoxic-ischemic injury complicates inflicted and accidental traumatic brain injury in young children: the role of diffusion-weighted imaging J Neurotrauma. 2007 Jan;24:106–118.

3. Kweon HJ, Suh BC. Acid-sensing ion channels (ASICs): therapeutic targets for neurological diseases and their regulation BMB Rep. 2013 Jun;46:295–304.

4. Balestrino M, Somjen GG. Concentration of carbon dioxide, interstitial pH and synaptic transmission in hippocampal formation of the rat J Physiol. 1988 Feb;396:247–266.

5. Caspers H, Speckmann EJ. Cerebral pO 2, pCO 2 and pH: changes during convulsive activity and their significance for spontaneous arrest of seizures Epilepsia. 1972 Sep;13:699–725.

6. Yang XF, Shi XY, Ju J, Zhang WN, Liu YJ, Li XY, et al. 5% CO_2_ inhalation suppresses hyperventilation-induced absence seizures in children Epilepsy Res. 2014 Feb;108:345–348.

7. Woodbury DM, Karler R. The role of carbon dioxide in the nervous system Anesthesiology. 1960 Nov-Dec;21:686–703.

8. Lennox W. The effect on epileptic seizures of varying the composition of the respired air. Journal of Clinical Investigation. 1928;6:23–24.

9. Tolner EA, Hochman DW, Hassinen P, Otahal J, Gaily E, Haglund MM, et al. Five percent CO(2) is a potent, fast-acting inhalation anticonvulsant Epilepsia. 2011 Jan;52:104–114.

10. Yang XF, Shi XY, Ju J, Zhang WN, Liu YJ, Li XY, et al. 5% CO(2) inhalation suppresses hyperventilation-induced absence seizures in children Epilepsy Res. 2014 Feb;108:345–348.

11. Costine-Bartell BA, McGuone D, Price G, Crawford E, Keeley KL, Pareja JM-, et al. Development of a Model of Unilateral Hemispheric Hypodensity (“Big Black Brain”) Journal of Neurotrauma. 2019;36:815–833.

12. Foster KA, Recker MJ, Lee PS, Bell MJ, Tyler-Kabara EC. Factors associated with hemispheric hypodensity after subdural hematoma following abusive head trauma in children J Neurotrauma. 2014 Oct 1;31:1625–1631.

13. Dias MS, Backstrom J, Falk M, Li V. Serial radiography in the infant shaken impact syndrome Pediatr Neurosurg. 1998 Aug;29:77–85.

14. Gilles EE, Nelson MD, Jr. Cerebral complications of nonaccidental head injury in childhood Pediatr Neurol. 1998 Aug;19:119–128.

15. Khan NR, Fraser BD, Nguyen V, Moore K, Boop S, Vaughn BN, et al. Pediatric abusive head trauma and stroke J Neurosurg Pediatr. 2017 Aug;20:183–190.

16. Costine-Bartell BA, McGuone D, Price G, Crawford E, Keeley KL, Munoz-Pareja J, et al. Development of a Model of Hemispheric Hypodensity (“Big Black Brain”) J Neurotrauma.2019 Mar 1;36:815–833.

17. Costine-Bartell B, Price G, Shen J, McGuone D, Staley K, Duhaime AC. A perfect storm: The distribution of tissue damage depends on seizure duration, hemorrhage, and developmental stage in a gyrencephalic, multi-factorial, severe traumatic brain injury model Neurobiol Dis. 2021 Mar 19:105334.

18. Missios S, Harris BT, Dodge CP, Simoni MK, Costine BA, Lee YL, et al. Scaled cortical impact in immature swine: effect of age and gender on lesion volume J Neurotrauma. 2009 Nov;26:1943–1951.

19. Missios S, Harris BT, Dodge CP, Simoni MK, Costine BA, Lee YL, et al. Scaled cortical impact in immature swine: effect of age and gender on lesion volume J Neurotrauma. 2009 Nov;26:1943–1951.

20. Costine-Bartell B, Price G, Shen J, McGuone D, Staley K, Duhaime AC. A perfect storm: The distribution of tissue damage depends on seizure duration, hemorrhage, and developmental stage in a gyrencephalic, multi-factorial, severe traumatic brain injury model Neurobiol Dis. 2021 Jul;154:105334.

21. Dulla CG, Dobelis P, Pearson T, Frenguelli BG, Staley KJ, Masino SA. Adenosine and ATP link PCO2 to cortical excitability via pH Neuron. 2005 Dec 22;48:1011–1023.

22. Ziemann AE, Schnizler MK, Albert GW, Severson MA, Howard MA, 3rd, Welsh MJ, et al. Seizure termination by acidosis depends on ASIC1a Nat Neurosci. 2008 Jul;11:816–822.

23. Chesler M. Regulation and modulation of pH in the brain Physiol Rev. 2003 Oct;83:1183–1221.

24. Ziemann M, Sedemund-Adib B, Reiland P, Schmucker P, Hennig H. Increased mortality in long-term intensive care patients with active cytomegalovirus infection Crit Care Med. 2008 Dec;36:3145–3150.

25. Rivera C, Wegelius K, Reeben M, Kaila K, Michael P. Different sensitivities of human and rat rho(1) GABA receptors to extracellular pH Neuropharmacology. 2000 Apr 3;39:977–989.

26. Huang RQ, Dillon GH. Effect of extracellular pH on GABA-activated current in rat recombinant receptors and thin hypothalamic slices J Neurophysiol. 1999 Sep;82:1233–1243.

27. Guaranha MS, Garzon E, Buchpiguel CA, Tazima S, Yacubian EM, Sakamoto AC. Hyperventilation revisited: physiological effects and efficacy on focal seizure activation in the era of video-EEG monitoring Epilepsia. 2005 Jan;46:69–75.

28. Xiong ZQ, Saggau P, Stringer JL. Activity-dependent intracellular acidification correlates with the duration of seizure activity J Neurosci. 2000 Feb 15;20:1290–1296.

29. Tamim I, Chung DY, de Morais AL, Loonen ICM, Qin T, Misra A, et al. Spreading depression as an innate antiseizure mechanism Nat Commun. 2021 Apr 13;12:2206.

30. Hartings JA, York J, Carroll CP, Hinzman JM, Mahoney E, Krueger B, et al. Subarachnoid blood acutely induces spreading depolarizations and early cortical infarction Brain. 2017 Oct 1;140:2673–2690.

31. Sugimoto K, Yang J, Fischer P, Takizawa T, Mulder IA, Qin T, et al. Optogenetic Spreading Depolarizations Do Not Worsen Acute Ischemic Stroke Outcome Stroke. 2023 Apr;54:1110–1119.

32. Costine-Bartell BA, Declan McGuone, George Price, Eleanor Crawford, Kristen L. Keeley, Jennifer Munoz-Pareja, Carter P. Dodge, Kevin Staley, and Ann-Christine Duhaime. Development of a Model of Unilateral Hemispheric Hypodensity Journal of Neurotrauma. 2019:815–833.

33. Duhaime AC, Durham S. Traumatic brain injury in infants: the phenomenon of subdural hemorrhage with hemispheric hypodensity (“Big Black Brain”) Prog Brain Res. 2007;161:293–302.

34. White A, Williams PA, Hellier JL, Clark S, Dudek FE, Staley KJ. EEG spike activity precedes epilepsy after kainate-induced status epilepticus Epilepsia. 2010 Mar;51:371–383.

35. Perucca P, Dubeau F, Gotman J. Intracranial electroencephalographic seizure-onset patterns: effect of underlying pathology Brain. 2014 Jan;137:183–196.

